# Methyl-CODEC enables simultaneous methylation and duplex sequencing

**DOI:** 10.1101/2024.12.23.630102

**Authors:** Ruolin Liu, Farzaneh Darbeheshti, Laurel Walsh, Rachel Li, Jin H. Bae, Hayet Radia Zeggar, Azeet Narayan, Kan Xiong, G. Mike Makrigiorgos, Viktor A. Adalsteinsson

## Abstract

DNA mutations and methylation often contribute to disease development in a synergistic manner. While duplex sequencing is the most accurate method for detecting DNA mutations, it typically lacks the ability to simultaneously assess methylation or requires many reads. Here, we developed Methyl-CODEC to enable simultaneous methylation sequencing and duplex sequencing using single read pairs. To achieve this, Methyl-CODEC links an enzymatically deaminated sense strand to the reverse complement of the antisense strand, which is protected from conversion by using conversion-resistant dCTPs in the strand linking step of CODEC. Methyl-CODEC shows high concordance with standard enzymatic or bisulfite based whole genome methylation sequencing, while also uniquely preserving the original DNA sequence. This improves genetic sequencing accuracy, enables better alignment for next-generation sequencing (NGS), and distinguishes C>T mutations from unmethylated Cs. It also identifies rare mutations including those producing methylated Cs, which are enriched in CpG contexts. Methyl-CODEC opens new horizons for enhanced detection of biomarkers in cancer and molecular medicine.

## Introduction

DNA molecules carry not only genetic information within their canonical nucleobases but also crucial epigenetic modifications. Together, these genetic and epigenetic alterations influence cellular behavior in a synergistic manner, and a comprehensive assessment of both is essential for effective disease detection and treatment. While duplex sequencing^1^ is considered the gold standard for highly accurate genetic sequencing, high numbers of reads are often required when both strands are sequenced separately. To overcome this issue, we recently developed CODEC^2^ which links both strands of each DNA duplex together and enables them to be sequenced together. For epigenetic analyses, methods such as Whole Genome Bisulfite Sequencing (WGBS)^3^ and Enzymatic Methyl Sequencing (EM-seq)^4^ are widely utilized. Both methods convert unmethylated cytosine (C), which is the predominant state of cytosine in the genome (> 95%), into uracil (U), and subsequently into thymine (T). This process results in requiring a ‘three-letter’ genome for read alignment, which can hinder alignment accuracy and efficiency. Recently, TET-assisted pyridine borane sequencing (TAPS)^5^ and direct methylation sequencing (DM-seq)^6^ have been introduced, offering improved mapping due to selectively converting 5-methylcytosine (5mC) instead of unmethylated C. However, a significant limitation of all these methods is their difficulty in distinguishing C to T conversion via enzymatic or chemical deamination from C>T mutation, which is the most prevalent mutation in nature^7,8^.

Other technologies capable of simultaneous epigenetic and genetic sequencing have been developed. Third-generation sequencing technologies, such as Oxford Nanopore Technologies (ONT)^9,10^ and Pacific Biosciences (PacBio)^11^, enable direct readouts of both epigenetic and genetic sequences. However, their genetic sequencing accuracy is typically lower than that of short-read sequencing and falls far below the accuracy of duplex sequencing, except when consensus reads are utilized, but this compromises throughput. Additionally, they often require high input amounts of unamplified DNA, which constrains their applicability to samples with limited DNA, such as liquid biopsies.

For short-read sequencing, methods for simultaneous epigenetic and genetic sequencing have been developed that make a copy of single-stranded DNA and use the original strand for methylation detection and the copied strand for mutation detection. Methods like Five-letter sequencing^12^ and Methyl-SNP-seq^13^ allow simultaneous sequencing of epigenetic and genetic data in a single workflow, but their detection of rare mutations is limited by single-strand accuracy (error rate ∼10^-4^). MethylSaferSeqS^14^ copies both strands of a DNA duplex using a primer that is biotinylated to permit the separation of the original and copied strands. This enables methylation sequencing of the original strands as well as Duplex Sequencing of the copied strands and has shown promising potential in the analysis of hundreds of patient plasma samples. However, it produces two separate libraries and requires sequencing each strand on its own. For assessing regional aggregated methylation levels (RAMLs) alongside genome sequencing in a single workflow, RIMS-seq2^15^ employs limited deamination to achieve 1% C to T transitions at 5mC sites. Yet, it does not provide single-base methylation resolution and has significant limitations in detecting rare somatic C>T mutations. At present, methods for simultaneous epigenetic and genetic sequencing have limitations in accuracy, efficiency, or breadth.

Here, we introduce Methyl-CODEC for simultaneous methylation and duplex sequencing using single read pairs. Based on our previous method, CODEC, which links both strands of each DNA duplex to achieve 1000-fold higher accuracy than standard NGS, Methyl-CODEC links an enzymatically methyl-converted sense strand with the copied antisense strand of the native sequence, and vice versa. This retains the duplex sequencing accuracy of CODEC while providing uncompromised alignment and single-base methylation resolution, allowing for simultaneous, genome-wide detection of methylation and mutations in single DNA duplexes using single read pairs.

## Results

### Methyl-CODEC workflow

Methyl-CODEC enables simultaneous epigenetic and genetic sequencing with the accuracy of duplex sequencing by linking a methyl-converted sense strand with the reverse complement of the antisense strand of the native sequence, and vice versa. This is achieved by using conversion-resistant dCTP analogs in place of standard dCTPs during the strand displacement step of CODEC^2^, followed by EM-seq conversion^4^ (Fig. 1a and Supplementary Fig. 1a). Specifically, after end repair and ligation to the CODEC adaptor quadruplex—in which standard dCTPs are replaced by conversion-resistant dCTPs—strand displacing extension occurs using a mix of conversion-resistant dCTP and regular dATP, dGTP, and dTTP. These conversion-resistant dCTPs are protected from being converted to Us (and subsequently to Ts during PCR) by APOBEC (Supplementary Fig. 1b). This process allows the copied side of each original strand to remain intact after methylation conversion (referred to as the protected strand, in contrast to the original/converted strand). From each input DNA duplex, Methyl-CODEC creates two products: one with the sense strand converted and the other with the antisense strand converted (Fig. 1a). This retains complete epigenetic and genetic information from both strands in the library.

**Fig. 1.**
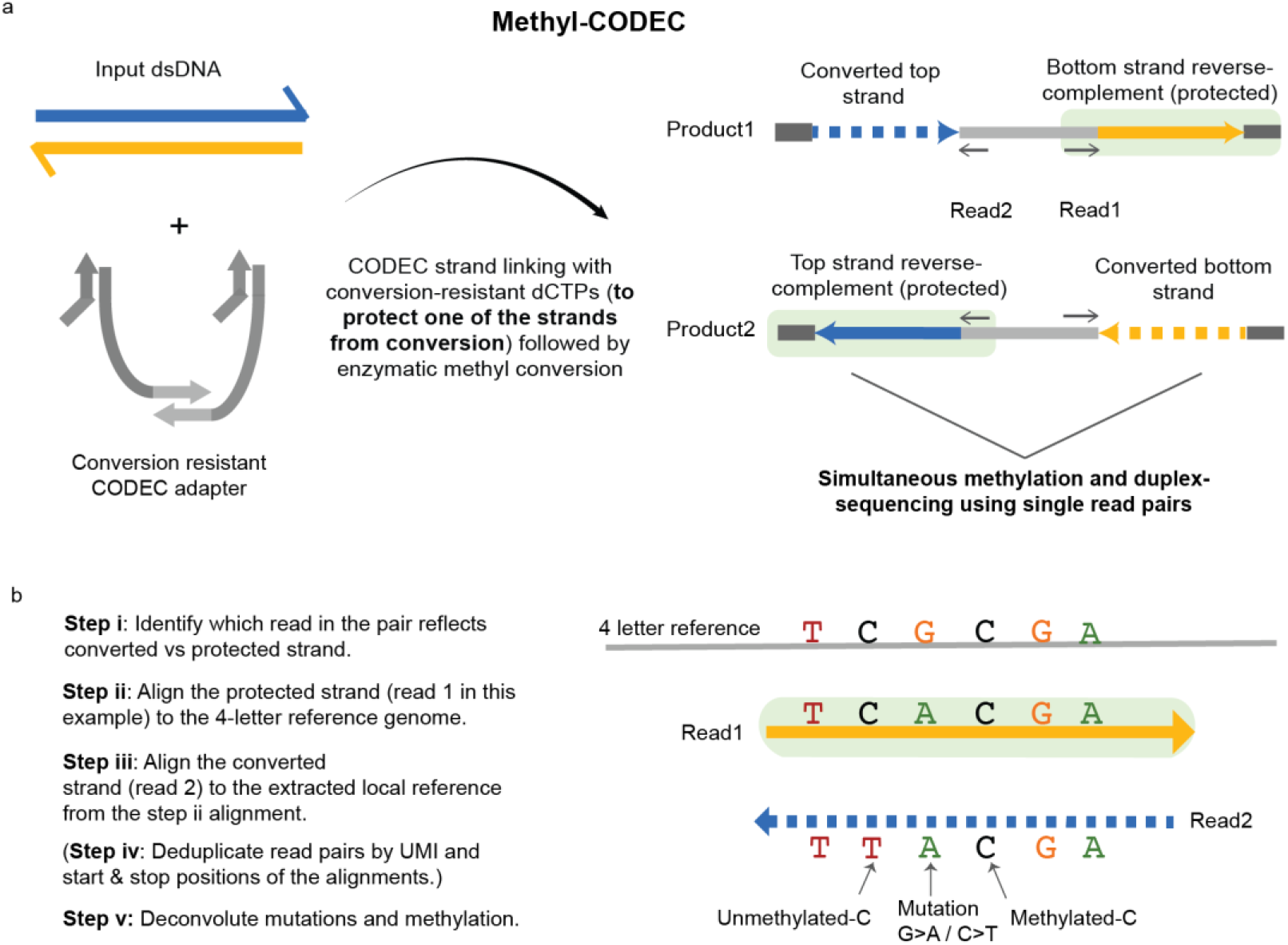
Methyl-CODEC library preparation and analysis workflows. **a**, Methyl-CODEC captures both genetic and epigenetic information in a single workflow. A modified CODEC adapter quadruplex, which is resistant to conversion, is used for ligation. The strand-displacing extension is performed using Phi29 polymerase with conversion-resistant dCTPs, instead of regular dCTPs, followed by enzymatic methyl-conversion. This results in two products, each comprising the converted sequence of one strand (dashed line) concatenated with the reverse complement of the other strand with conversion protection (solid line with green background). Unique molecular identifiers (UMIs) are included in the adapters to allow for the two products to be re-associated as needed, although a single Methyl-CODEC product is sufficient to achieve duplex accuracy. **b**, Methyl-CODEC analysis workflow. i) The strand information (protected vs. converted) is determined for a given read based on G% or C/T or G/A mismatches. ii) For a given duplex read pair, the protected strand is first aligned to the standard 4-letter reference genome (HG38) by BWA-MEM. The local reference sequence is extracted from the alignment of the protected strand. iii) The converted strand is aligned to the extracted local reference sequence by BWA-SW using a custom setting that does not penalize C/T or A/G mismatches. iv) PCR duplicates are marked using UMIs, start and stop positions of the aligned paired-end reads, and only the fragment with the best sum of quality scores was used for down-stream analysis. v) A,G,T, unmethylated C and methylated C are determined using the duplex informed deconvolution table (Supplementary Fig. 1c) from comparing the sequences of the two strands. Mutations are called by comparing to the reference. Owing to the use of complementary strands, Methyl-CODEC achieves duplex accuracy for mutation detection. See Methods for further details.

A correct Methyl-CODEC product has one read of a pair derived from the protected strand and the other read from the converted strand, originating from opposing strands of the same original DNA duplex (i.e., duplex reads) (Fig. 1a), whereas Methyl-CODEC byproducts are generated in similar ways as CODEC^2^ byproducts. Moving forward, we will refer to the correct Methyl-CODEC product simply as “Methyl-CODEC product” to distinguish it from Methyl-CODEC byproducts. To determine Methyl-CODEC product reads, the protected strands are first aligned to the entire reference genome using BWA-MEM^16^. Next, the converted strands are aligned strictly to regions where protected strands were aligned, with an additional 500 bp padding, using custom BWA-SW^16^ with 0 mismatch penalty for conversion induced C/T or G/A mismatches (Fig. 1b & Methods). Only reads with a minimum alignment score of 30 for both strands are considered Methyl-CODEC products; other fragments are classified as byproducts but retained for further analysis. From deduplicated Methyl-CODEC products, A, C, G, T and 5mC calls are made based on the deconvolution table (Supplementary Fig. 1c) in overlapping regions of the two strands and subject to fragment level and base level filtering, similar to CODEC (see Methods). Methylation calls are stored as Bismark-like tags^17^ for compatibility of downstream analysis. We first confirmed that the Methyl-CODEC workflow can generate Methyl-CODEC products using the human cell line NA12878. Using 4 different conversion-resistant analogs (5m-dCTP, carboxy-dCTP, hydroxy-dCTP and propargylamino-dCTP), 47.6%, 76.6%, 71.2% and 76.8% of the total reads, respectively, were confirmed as Methyl-CODEC products, with the majority (∼95%) of Methyl-CODEC byproducts being intermolecular byproducts (Supplementary Table 2). Overall, hydroxy-dCTP was selected as the conversion-resistant dCTP variant for Methyl-CODEC due to its most complete protection against enzymatic methyl-conversion (Supplementary Fig. 3).

### Methyl-CODEC matches EM-seq WGMS and WGBS for methylation sequencing

Methyl-CODEC utilizes the same enzymatic methyl-conversion step as EM-seq^4^, while WGBS remains a widely used approach for methylation analysis. Therefore, we compared methylation results from whole genome Methyl-CODEC sequencing with Whole Genome Methylation Sequencing (WGMS) data from EM-seq and WGBS on the same cell line, NA12878, obtained from public repositories^4^. For alignment of EM-seq and WGBS data, Bismark^17^ was used. First, we found that genome-wide average cytosine methylation levels across three different contexts were similar (Fig. 2a). Additionally, the per-CpG island methylation level from Methyl-CODEC showed high concordance with EM-seq and WGBS, achieving Pearson correlations of 0.99 and 0.97, and strong linear fits with adjusted R^2^ of 0.97 and 0.95, respectively (Fig. 2b).

**Fig. 2.**
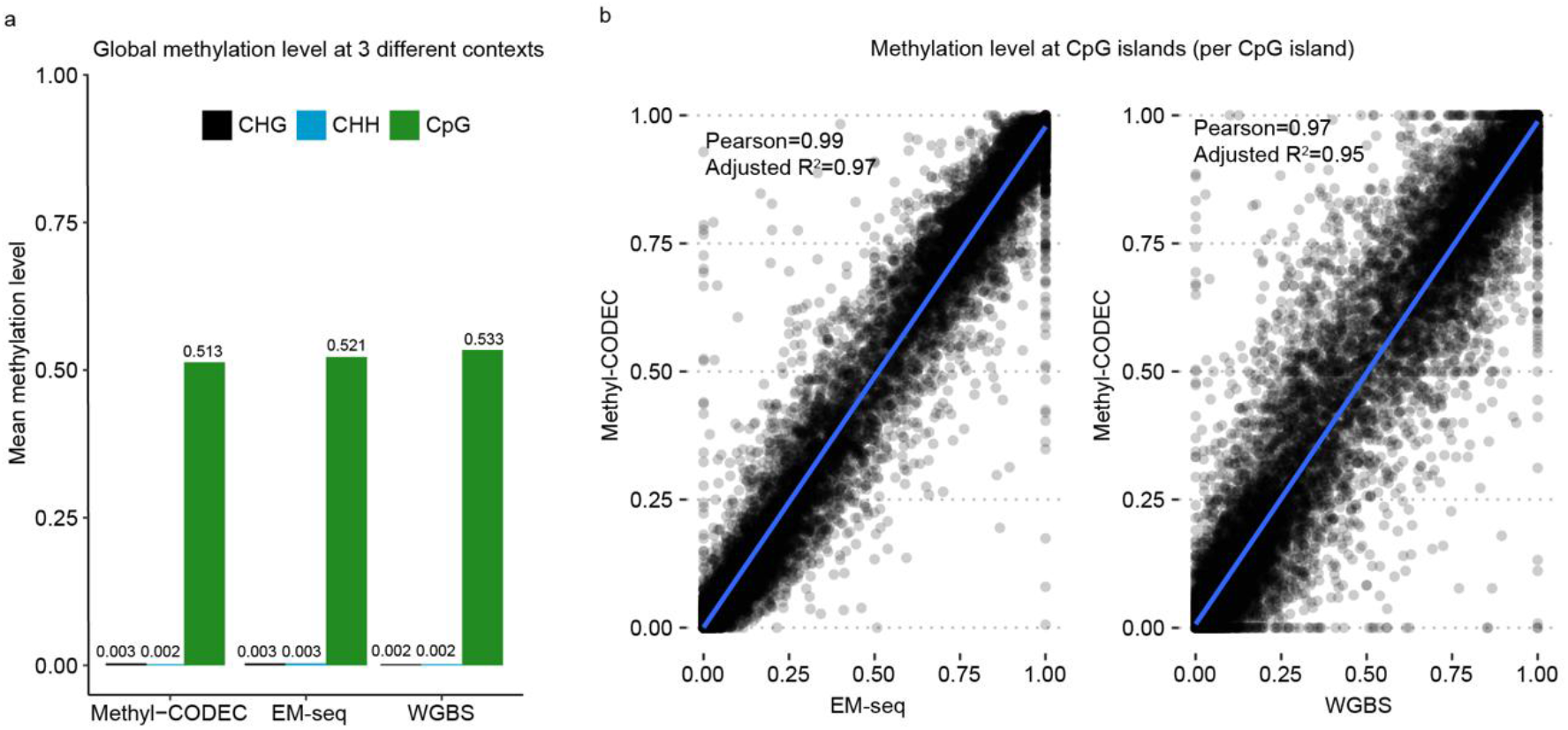
Concordance of 5-methylcytosine readout between Methyl-CODEC and EM-seq or WGBS. **a**, Genome-wide methylation level at three different sequence contexts (CpG, CHG, and CHH). H is an IUPAC code which stands for A, C, or T. **b**, Scatter plots comparing methylation level per CpG island (n=25,368) between Methyl-CODEC and EM-seq, and between Methyl-CODEC and WGBS. The blue lines show linear fits made using R ggplot2. Pearson correlations and adjusted R-squares are shown in the top left corners.

Unlike EM-seq and WGBS, Methyl-CODEC uniquely preserves the original DNA sequence for NGS alignment. To assess the impact of this on read alignment, we first removed duplicates and then simulated paired-end WGMS from NA12878 Methyl-CODEC and aligned them by Bismark (see Methods). We compared the concordance between the Methyl-CODEC alignments based on the original 4-letter DNA sequence, which is how Methyl-CODEC is configured, and the Bismark alignment based on the methyl-converted sequence (3-letters) as per WGBS or EM-seq. After applying mapping quality thresholds for both Methyl-CODEC and Bismark, we observed a sharp difference when using different Bismark alignment parameters (Supplementary Fig. 2a). In particular, Bismark’s local alignment mode significantly reduced the number of unaligned reads compared to the default mode (11.3% vs 28.9% unaligned). However, this improvement came with a slight increase in the fraction of misplaced/discordant reads: 0.38% in local mode compared to 0.06% in default mode (Supplementary Fig. 2a). Because Methyl-CODEC does not rely upon a ‘3-letter’ genome for read alignment, our results suggest that Methyl-CODEC may enable more accurate read alignment for methylation sequencing, although its impact was limited to a minor subset of reads with likely incorrect alignments.

In addition, Methyl-CODEC can distinguish C>T mutations, including germline and somatic or rare mutations, from C to T conversions via enzymatic or chemical deamination, given its retention of the original DNA sequence. To explore this, we prepared and sequenced 3 additional Methyl-CODEC libraries using Human HCT116 DKO samples: fully unmethylated DNA, fully methylated DNA, and a 1:19 mix of these two. From the parent HCT116 cell line^18^, we detected 590,281 germline mutations, including indels, at CpG sites, with about 80% of these germline mutations being C>T/G>A homozygous and heterozygous SNPs (Supplementary Fig. 2b). We hypothesized that for EM-seq and WGBS, this could potentially result in false calling of hundreds of thousands of unmethylated CpG sites, which could lead to an appreciable underestimation of global methylation. Indeed, using the fully methylated genomes, we observed 97.3% global CpG methylation with Methyl-CODEC, compared to 95.8% (Bismark-default) and 92.4% (Bismark-local) global CpG methylation with simulated paired-end EM-seq WGMS (Supplementary Fig. 2c). We then calculated CpG methylation levels both outside and within the germline C>T/G>A sites. Methyl-CODEC consistently maintained global methylation levels in both cases (with 97.3% outside the germline C>T/G>A sites and 97.4% within these sites, closely aligning with the global average of 97.3%). In contrast, simulated EM-seq WGMS aligned by Bismark-default achieved a similar methylation level (97.0%) to Methyl-CODEC outside the germline C>T/G>A regions (Supplementary Fig. 2d) but dropped significantly to 31.6% at germline C>T/G>A sites (Supplementary Fig. 2e). Of note, this shortfall of EM-seq WGMS could be overcome by also performing whole-genome sequencing on the same sample, but this introduces added complexity. For the fully unmethylated genome, which contains low levels of DNA methylation^19^, all methods showed similar CpG methylation levels at 13.4%, 13.4%, and 13.1%, respectively. In all, Methyl-CODEC is superior to EM-seq and WGBS in distinguishing C>T mutations from C to T conversions via enzymatic or chemical deamination.

### Methyl-CODEC enables duplex sequencing with similarly high accuracy as CODEC

Duplex sequencing ^1^, as well as CODEC, rely on the principle that sequence information obtained from both DNA strands can distinguish genuine mutations from sequencing noise^20^. To determine whether duplex sequencing accuracy is retained for Methyl-CODEC, we analyzed the NA12878 cell line for mutation detection (see Methods). This analysis yielded a raw deduplicated depth of 25x (duplex depth of 5x), with 41% of duplexes covered by just single read pairs of Methyl-CODEC products. Our results confirmed that Methyl-CODEC products have the required accuracy across all base contexts, including C>T/G>A mutations mapped to the ‘G’ reference bases (Fig. 3a). This is achieved because the G nucleotide on the strand complementary to a C (referred to as the ‘G-strand’ in Bis-SNP^21^) remains unaffected by enzymatic methyl-conversion. On the other hand, the C>T/G>A mutations mapped to ‘C’ reference are prone to false detection, possibly due to 1) PCR or sequencing errors and/or 2) incomplete protection (Fig. 3b). Additionally, we hypothesized that impurities in commercially available dCTP analogs can lead to incomplete protection from enzymatic conversion. Indeed, we found that hydroxy-dCTP gave the best C>T accuracy (a 2-log reduction in error rate compared to 5m-dCTP) when ‘C’ is the reference (Supplementary Fig. 3). The mutation frequency of C>T was close to the raw Illumina sequencing error rate (1x10^-3^), suggesting that incomplete protection was almost eliminated by using hydroxy-dCTP. On the other hand, propargylamino-dCTP yielded worse accuracy, and carboxy-dCTP yielded accuracy between that of 5m-dCTP and hydroxy-dCTP (Supplementary Fig. 3). In addition, T>C mutations were also prone to false detection by PCR or sequencing errors due to unmethylated C being indistinguishable from a single T>C sequencing error on the protected strand (Fig. 3b), and we did observe the T>C mutation rate was close to the raw Illumina sequencing error rate (Fig. 3a). For all single base contexts, except for C>T and T>C, the residual SNV frequencies exhibited minimal or no strand bias, indicating all Methyl-CODEC products can be used for their detection with duplex sequencing accuracy. Meanwhile, for C>T and T>C, duplex sequencing accuracy is still attainable when the complementary mutation (e.g., G>A or A>G) is detected, which will be present on one of two Methyl-CODEC products from each DNA duplex.

**Fig. 3:**
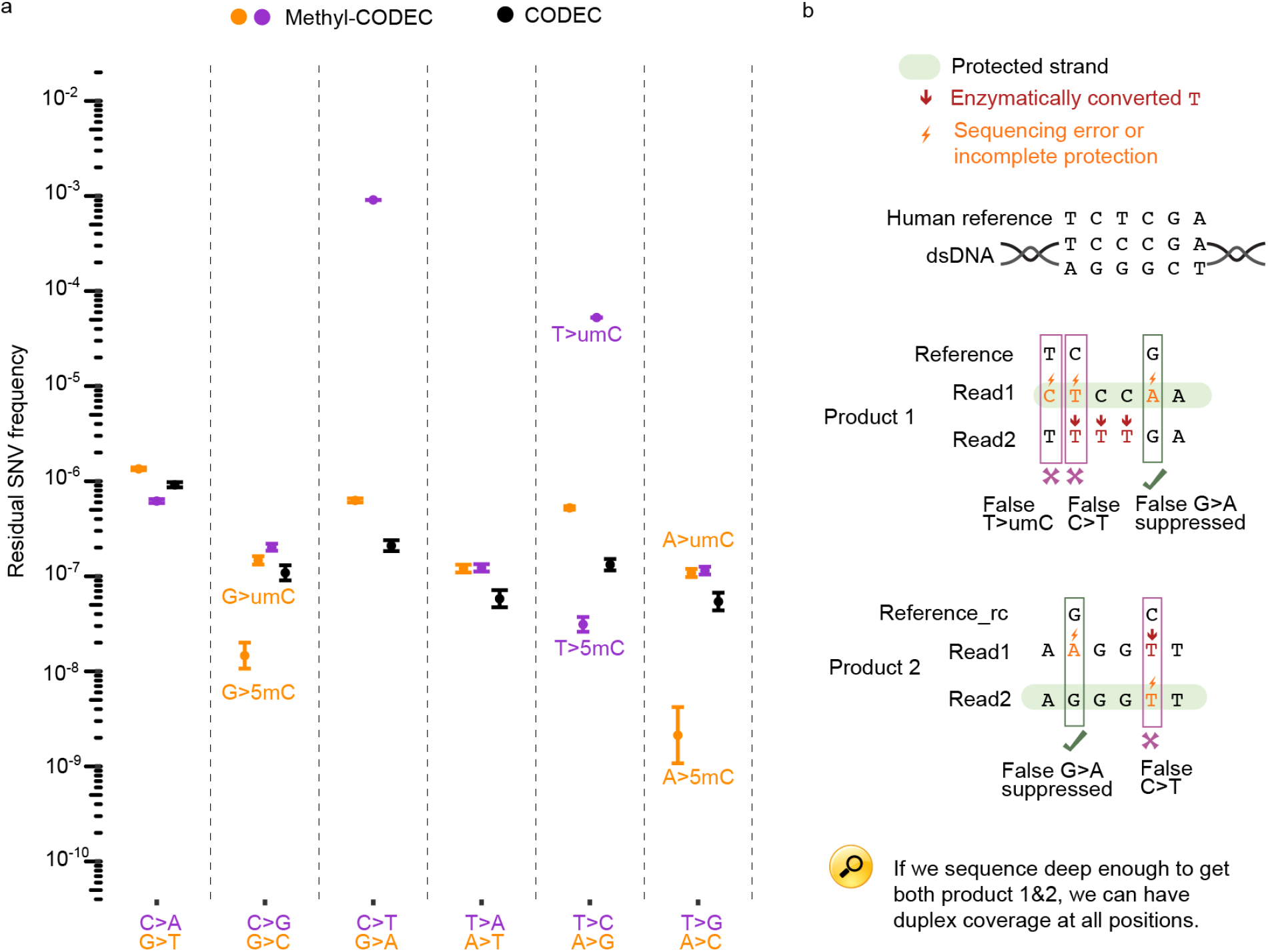
Methyl-CODEC achieves duplex sequencing level accuracy (> Q60). **a**, A scatter plot comparing residual SNV frequencies of Methyl-CODEC (raw deduplicated depth at 25x, duplex depth at 5x) and standard CODEC in NA12878, both using a commercial end-repair/dA-tailing protocol. Methyl-CODEC data is split into pyrimidines (C or T) as the reference and purines (G or A) as the reference, while CODEC data represents the average across both. Error bars indicate 95% binomial confidence intervals by Wilson method. Mutations that produce methylated cytosines (5mC) and unmethylated cytosines (umC) are labeled next to the corresponding dots. **b**, A hypothetical double stranded DNA example demonstrating how to use Methyl-CODEC strand information (product 1 vs product 2) to achieve duplex accuracy for C>T/G>A contexts and how T>umC can result from sequencing errors. “rc” indicates reverse complement.

### Methyl-CODEC expands genome-wide mutation calling

The conventional single base substitution (SBS) alphabet, e.g., G>C, considers only mutations among the standard bases A, C, G, and T for both reference and alternative alleles. Methyl-CODEC extends this by distinguishing C or cytosine into non-methylated cytosine (umC) and 5-methylcytosine (5mC) as distinct alternative alleles, denoted as A>umC, G>umC, T>umC and A>5mC, G>5mC, T>5mC respectively (Fig. 3a). In NA12878 Methyl-CODEC data, we found that 5 out of the 6 expanded mutation types have residual mutation frequencies close to or lower than those of the corresponding conventional mutation types in CODEC (Fig. 3a), with T>umC being an exception, which we reason was dominated by sequencing errors on the protected strand (Fig. 3b), but could be called with duplex accuracy if the other Methyl-CODEC product were sequenced. On the other hand, T>umC cannot be detected by standard enzymatic or bisulfite based methylation sequencing, as it is impossible to distinguish between naturally occurring T and T generated from enzymatic or chemical conversion of unmethylated C. Interestingly, observed frequencies for A>5mC, G>5mC and T>5mC were significantly lower (between 10^-9^ and 10^-7^), aligning with our expectations, as 5mC predominantly occurs at CpG dinucleotides which comprise only 1% of the genome.

We then investigated the proportion of rare duplex mutations in NA12878 Methyl-CODEC that resulted in the formation of novel cytosine (7.8%) and CpG dinucleotides (2.3%, Fig. 4a), after removing C>T and T>umC due to their sub-duplex-sequencing accuracy. Next, we analyzed the trinucleotide contexts of the 148 rare duplex mutations creating 5mC and the 656 rare duplex mutations creating umC, and observed that the majority (82%) of the novel 5mC mutations occurred at newly formed CpG sites, particularly for T>5mC and A>5mC, whereas only 17% of the novel umC mutations formed at CpG sites (Fig. 4b), consistent with established data and models indicating that methylation predominantly occurs at CpG sites in humans. Interestingly, for G>5mC mutations, a significant portion (48.6%) did not occur at newly formed CpG sites. As methylation is assessed from only a single strand of each original DNA duplex using a single Methyl-CODEC product, it is possible that methylation accuracy could be further improved if both Methyl-CODEC products from each duplex were recovered. Nonetheless, our results suggest the feasibility of detecting rare mutations and their corresponding methylation status in single DNA duplexes, using single NGS read pairs, with Methyl-CODEC.

**Fig. 4:**
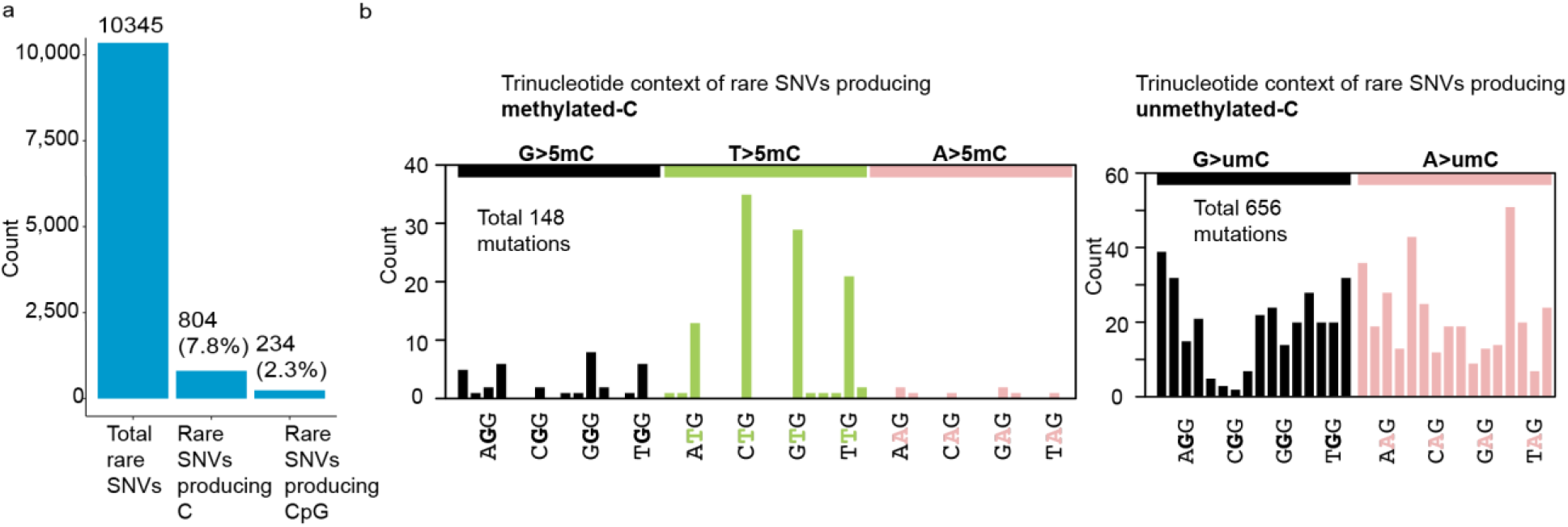
Expanded mutation calling of Methyl-CODEC in NA12878. a) A bar plot showing, from left to right, the total number of rare duplex SNVs, the number of rare duplex SNVs which produced a C, and the number of rare duplex SNVs which created new CpG sites, excluding C>T and T>umC which are not at duplex accuracy. b) Plots showing the count of rare duplex SNVs which produced a 5mC (left) vs umC (right) across trinucleotide contexts. Reference trinucleotide contexts which can generate novel CpG are indicated on the X-axis. T>umC is excluded due to lack of duplex accuracy.

## Discussion

Methyl-CODEC is a simple yet powerful adaptation of CODEC that enables simultaneous methylation and duplex sequencing using single read pairs. It introduces conversion-resistant adapters and dCTP analogs during CODEC strand linking and adds an enzymatic methyl conversion step along with a novel analysis workflow. While prior methods have been developed for combined genetic and epigenetic analysis^12–15^, Methyl-CODEC is the first to achieve both genetic duplex sequencing accuracy and single-base methylation resolution using single read pairs at the whole genome level.

To demonstrate this, we compared Methyl-CODEC’s genetic sequencing accuracy to standard CODEC and its methylation sequencing performance to EM-seq WGMS and WGBS on the same cell line. Our results showed that Methyl-CODEC can achieve duplex sequencing accuracy for all single nucleotide contexts using single read pairs, including for C>T and T>C when G>A and A>G, respectively, are instead evaluated. For methylation sequencing, Methyl-CODEC demonstrated strong concordance with widely used techniques such as WGBS or EM-seq, while uniquely preserving the original DNA sequence for improved NGS alignment and C>T mutation detection. We found that Methyl-CODEC uniquely improved methylation detection at CpG sites (around 590,000 sites) overlapping germline C>T/G>A mutations. Methyl-CODEC also expanded genome-wide mutation calling by distinguishing C into umC and 5mC as distinct alternative alleles. This enabled us to discover genome-wide, low-frequency mutations and methylation co-occurring within single base pairs of single DNA duplexes. Those novel 5mC were found to be enriched in newly formed CpG contexts compared to non CpG contexts, suggesting that most of these novel 5mC sites were genuine and may reveal new biological processes. We believe this could be useful in many areas such as for cancer and minimal residual disease detection using liquid biopsies.

A current limitation of Methyl-CODEC in epigenomic sequencing is its inability to distinguish between 5mC and 5hmC, although this could be added in the future. In addition, the current end-repair/dA-tailing step leads to artificial hypomethylation in the resynthesized regions, although we among others have described superior end-repair/dA-tailing methods that could limit resynthesis^22,23^. Furthermore, only about half of the coverage is usable for detecting C>T and T>C at duplex accuracy given the need to detect these by way of G>A and A>G, respectively. Although Methyl-CODEC generates two Methyl-CODEC products per original DNA duplex with each of the respective base changes, recovering both in a WGS library is rare. Methyl-CODEC, like CODEC, is conceptually compatible with targeted sequencing workflows. This has several conceptual advantages. For instance, by preserving a native strand sequence, Methyl-CODEC hybrid capture sequencing stands to streamline and mitigate potential biases typically associated with hybrid capture following bisulfite or enzymatic methyl-conversion.

Methyl-CODEC has the potential to enable a wide range of clinical applications. In oncology, for example, it can integrate tumor-specific epigenetic and genetic markers to enhance comprehensive cancer detection and management, thereby advancing precision oncology. Duplex sequencing accuracy is pivotal for sensitive detection of minimal residual disease from blood^24,25^ and Methyl-CODEC uniquely preserves this accuracy. By capturing epigenetic and genetic modifications in cell-free DNA (cfDNA), Methyl-CODEC could improve liquid biopsy-based cancer screening ^26–29^. For example, while mutations on cfDNA can indicate the presence of cancer^30^, epigenetic patterns detected on the same molecules can help identify the tissue of origin^31^. This capability can guide targeted imaging or biopsies to confirm the presence and extent of cancer prior to initiating therapy. Additionally, Methyl-CODEC can aid in the classification and selection of treatment strategies for cancers of unknown primary (CUP), a particularly challenging group to treat, by combining mutation with methylation anaysis^32,33^.

Beyond oncology, Methyl-CODEC could have applications in prenatal testing and transplantation, where it can identify the tissue of origin of DNA molecules derived from the fetus or tissue donor^34^. Methyl-CODEC could also be applied to other biological areas, such as detecting allele-specific methylation which is crucial for localizing regulatory sequence polymorphisms^35,36^ and assessing biological aging^37,38^. A recent study has shown that epigenetic clocks are influenced by somatic mutations, which can alter methylation patterns not only at the mutation site but also in surrounding regions^39^. Lastly, as Methyl-CODEC does not depend on a reference genome to derive methylation status, we envision applications in infectious diseases or in non-model organisms.

In conclusion, by enabling simultaneous methylation and duplex sequencing using single read pairs, Methyl-CODEC opens promising new avenues of broad potential impact for biomedicine.

## Methods

### Samples and oligonucleotides

Universal methylated and non-methylated standard DNA of HCT 116 DKO samples were purchased from ZYMO Research (Catalog no. D5014). Genomic DNA of the NA12878 cell line was obtained from Coriell Institute of Medical Research (Catalog no. GM12878). DNA fragmentation was performed by a Covaris ultrasonicator to achieve a mean fragment size of 150 bp. The fragmented DNA was purified via AMPure (Beckman Coulter), followed by elution in low TE buffer. 20ng of fragmented DNA was used as input for library preparation. To prepare a 95% non-methylated library, we mixed fragmented non-methylated standard DNA with fragmented methylated standard DNA in a 19:1 ratio. Methyl-CODEC adapter oligonucleotides containing methylated cytosine, instead of unmodified cytosine, were ordered from Integrated DNA Technologies (IDT) with polyacrylamide gel electrophoresis (PAGE) purification. Three different sets of adapter oligonucleotides with distinct indices were purchased (Supplementary Table 1). The Methyl-CODEC adapter quadruplex was formed by adding 5 μM of each oligonucleotide to low TE buffer with 100 mM NaCl in a total volume of 40 μl. The mixture was then heated at 85 °C for 3 min, gradually cooled to 20 °C, and incubated at room temperature for 12 h.

### Methyl-CODEC library preparation

#### End-repair/dA-tailing and ligation

NEBNext Ultra II DNA Library Prep Kit for Illumina (Catalog no. E7645) was used according to the manufacturer’s protocol (New England Biolab). 10-20 ng input DNA was eluted in low TE to a final volume of 50 μl. This was combined with 7 μl of End Prep Reaction Buffer and 3 μl of End Prep Enzyme Mix, followed by incubation at 20 °C for 30 min and then at 65 °C for another 30 min. Subsequently, DNA purification was performed using 1.8X AMPure XP beads according to the manufacturer’s protocol (Beckman Coulter), followed by elution in 60 μl of low TE buffer. Adaptor ligation was performed by adding 30 μl of Ligation Master Mix, 1 μl of Ligation Enhancer, and 3 μl of 5′-deadenylase (NEB, Catalog no. M0331S) to the 60 μl end-prepped product. Instead of using Illumina adaptors, we used 2.5 μl of 500 nM freshly diluted Methyl-CODEC quadruplex adaptor, and the incubation time was increased to 1 hour at 20°C. Purification was performed using 0.9x AMPure XP beads, followed by final elution in 40 μl of low TE buffer. Subsequently, 1.5 μl of USER (NEB, catalog no. M5505) was added, followed by incubation at 37 °C for 30 min.

#### Phi29 strand displacing extension

For the strand displacement step, Phi29 DNA polymerase (NEB, Catalog no. M0269L) was applied by adding the following to 41.5 μl of ligated DNA from the previous step, followed by incubation at 30 °C for 20 min: 10 μl of 10X buffer, 1 μl of polymerase, 47 μl of nuclease-free water, and 2 μl of 10 mM customized dNTPs, including dATP, dTTP, dGTP, and conversion-resistant dCTP analogues, evenly. We used four different conversion-resistant dCTPs, including 5-methyl-dCTP (NEB, Catalog no. N0356S), 5-Carboxy-dCTP (TriLink, Catalog no. N-2063), 5-Hydroxymethyl-dCTP (Jena Bioscience, Catalog no. NU-932S), and 5-Propargylamino-dCTP (Jena Bioscience, Catalog no. NU-809-1). The Phi29 extended product was purified using 0.75X AMPure XP beads, followed by final elution in 28 μl of nuclease-free water to prepare for the enzymatic methyl-conversion step.

#### Enzymatic methyl-conversion

NEBNext Methyl-seq Conversion Module (NEB, Catalog no. E7125S) was used for DNA conversion based on the manufacturer’s protocol, consisting of two main steps: TET2 Oxidation/BGT glucosylation and APOBEC Deamination. The former was done by adding the following to 28 μl Phi29 extended product: 10 μl TET2 buffer, 1 μl Oxidation Supplement, 1 μl Oxidation enhancer (BGT), 1 DTT, 4 μl TET2 enzyme, and 5 μl diluted (1:1250 ratio) FeII sulfate hexahydrate. The oxidation step occured at 37 °C for 1 h, followed by addition of 1 μl Stop Reagent and incubation at 37 °C for 30 min. Next, DNA was purified by 0.75X AMPure XP beads and final elution in 16 μl nuclease free water. The latter step, APOBEC Deamination, was initiated by DNA denaturation by adding 4 μl of formamide to 16 μl DNA and incubating at 85 °C for 10 min. Afterwards, the 20 μl denatured DNA was immediately transferred on ice followed by addition of 68 μl nuclease free water, 10 μl APOBEC buffer, 1 μl APOBEC, and 1 μl BSA and incubation for 3 h at 37 °C. Finally, the converted DNA was cleaned up by 0.75X AMPure XP beads and eluted in 16 μl nuclease free water.

#### Library amplification

Methyl-CODEC library amplification was performed by adding 25 μl Q5U Hot Start High-Fidelity DNA Polymerase (NEB, Catalog no. M0515) and 5 μl KAPA Library Amplification Primer Mix (Roche, Catalog no. KK2623) to 20 μl Methyl CODEC library molecules. PCR was run via Mastercycler X50 Eppendorf with the following program: Initial denaturation for 30 sec, then 8-10 cycles of 98 °C for 20 sec, 60 °C for 30 seconds, 65 °C for 2 min followed by final extension at 65 °C for 5 min. After PCR, the libraries were twice purified by 0.65X AMPure XP beads, consecutively, with 20 μl final elution. The libraries were quantified and qualified by Qubit dsDNA High Sensitivity kit (Invitrogen, Catalog no. Q33230) and Agilent High Sensitivity DNA Kit (Catalog no. 5067-4626).

### Sequencing

The 4 Methyl-CODEC NA12878 samples were sequenced using NovaSeq S4 with no index cycle and the read length of 169, whereas the 3 Methyl-CODEC HCT116 DKO samples were sequenced using NovaSeq X with 8 index cycles and the read length of 161. The 4 NA12878 samples had 930M (hydroxy-C), 166M (5mC), 999M (carboxy-C) and 419M (propargylamino-C) read pairs. The 3 HCT116 samples yielded 834M (methylated control), 820M (unmethylated control) and 772M read pairs (mixed). The sequencing metrics are available in the Supplementary Table 2.

### Methyl-CODEC analysis workflow

The Methyl-CODEC analysis workflow is based on CODEC workflow and shares many components. In brief, Methyl-CODEC uses the same demultiplexing and adapter trimming steps as CODEC.

i. **Confirming read structure**: After adapter trimming, Methyl-CODEC includes an additional step to determine whether a read pair has the correct structure (i.e., one read is from the converted strand, and the other is from the protected strand) and to identify its specific structure (i.e., product 1 vs product 2). This is achieved using a two-step algorithm. In the first step, if one read has a fraction of G less than 5% and the other has a fraction of G greater than 5%, the former is marked as the converted strand/read and latter as the protected strand/read. If both reads have a fraction of G either all above 5% or all below 5%, the second step is applied. In the second step, overlap alignment is performed, aligning the suffix of the first read to the prefix of the reverse complement of the second read using custom alignment parameters (e.g., no mismatch penalty for conversion induced C/T and G/A mismatches and high penalties for gaps). The C/T and G/A mismatches in the overlap region are counted. If the number of C/T mismatches is greater, the read pair is identified as product 1 (with the first read protected and the second converted). Conversely, if the number of G/A mismatches is larger, the read pair is identified as product 2 (with the first read converted and the second read protected). If no overlap alignment is found, the read pair is marked as a byproduct. The first step accelerates the process and accommodates cases where the two reads have little or no overlap.
ii. **Alignment of the protected reads**: Methyl-CODEC utilizes standard alignment algorithms without requiring methylation specific aligners. After confirming the read structure (i.e., product 1 vs product 2), Methyl-CODEC uses standard BWA-MEM^16^ to align the protected read to the entire reference genome (HG38 in this study). If the alignment is unsuccessful, the read-pair is marked as a byproduct. For successfully aligned reads, the reference sequences of the alignments are extracted, including 500 base pairs as padding.
iii. **Alignment of converted reads**. The converted reads are aligned to the local reference sequences obtained from the protected strand alignments using BWA-SW^16^. Custom alignment parameters are applied here as well, avoiding penalties for conversion-induced mismatches (i.e., C/T mismatches for product 1, and G/A mismatches for product 2). Reads with an alignment score below 30 are classified as byproducts.
iv. **Deduplication:** For all Methyl-CODEC product reads, Picard MarkDuplicates is used to mark PCR duplicates using UMIs, start and stop positions of the aligned paired-end reads. Among all PCR-duplicated read pairs of a single duplex, only the one with the best sum of quality scores was used for down-stream analysis in order to confirm that a single read pair is sufficient to achieve duplex accuracy. However, generating a consensus from duplicates may further improve accuracy.
v. **Determining methylation status and cytosine contexts**. After obtaining deduplicated paired-end alignments, the methylation statuses of cytosines in the overlapped regions are determined. For product 1, the Cs in the protected strands are first scanned. A methylated C or a non-methylated C is identified when the corresponding base on the converted strand is C or T, respectively, providing that the two complementary bases from each strand meet the quality score threshold (Q30). The same process applies to product 2 but focuses on G in the protected strand and G or A in the converted strand. In both situations, the 12 base pairs from the fragment ends are ignored. These methyl-resolved cytosines are then annotated based on their contexts (CpG, CHG or CHH, H is an IUPAC code which stands for A, C, or T) in a reference-independent manner. The context is called only if the adjacent bases from the two strands agree (accounting for conversion induced C/T, G/A mismatches) and either both meet the quality score threshold (Q30) or match the reference base. Cytosines in the CpG context are annotated with a Bismark-like symbol Z (methylated) or z (unmethylated). CHG and CHH contexts are determined based on the further adjacent bases. Finally all methyl-CODEC products with methylation status and context information are stored as a BAM file.

All steps (i-iv) are encapsulated in a single executable file. Alignments are performed in memory using direct calls to the BWA API in C++ code to minimize IO compute time.

### Methylation extraction

After generating BAM alignments with Bismark-like tags, fragment-level and locus-level methylation information was extracted using bismark_methylation_extractor, which also calculated sample-level methylation fractions for CpG, CHG, and CHH contexts. Methylation fractions for CpG islands were calculated by intersecting the locus-level methylation with CpG island coordinates.

### Processing EM-seq WGMS and WGBS

A 20-ng input EM-seq library and a 50-ng input WGBS library were downloaded from the EM-seq paper^4^. Reads were then aligned using Bismark (default mode), and methylation information was extracted with bismark_methylation_extractor, ignoring the first 12 bases from the read 2 (parameters: --ignore 2 --ignore_3prime 2 --ignore_r2 12 --ignore_3prime_r2 2) to adjust for M-bias^40^.

### Simulating paired-end EM-seq WGMS from Methyl-CODEC for comparison

Paired-end EM-seq WGMS data were simulated from Methyl-CODEC products by in-silico conversion of the protected strand (converting C to T for product 1 and G to A for product 2). The simulated paired-end WGMS data was aligned using Bismark in two modes (default and local). The local mode allows soft-clipping at the ends, while default mode enforces end-to-end alignment. One NA12878 and three HCT116 samples were simulated. In NA12878, the Methyl-CODEC alignments were compared with Bismark alignments. In three simulated HCT116 samples, after alignment by Bismark in two modes, methylation results were extracted from the original converted strands, excluding the first and last 12 bps in the reads, and then compared with Methyl-CODEC results.

### Extracting HCT116 methylation levels in CpG sites with and without overlapping germline mutations

Germline mutations in the HCT116 parental cell line were identified using GATK HaplotypeCaller^41^. Germline mutations (SNPs and Indels) overlapping CpG sites in the HG38 genome were further filtered to include only C>T and G>A SNPs. Methylation levels were calculated for CpG sites overlapping C>T and G>A SNPs and for CpGs that were unaffected by germline mutations, i.e., CpGs that were at least 3 bps away from any germline mutations.

### Somatic mutation detection

Methyl-CODEC uses the CODECsuite to call somatic mutations from single read pairs, with modifications to account for methylation conversion: i) Mutations are called if a base C from a protected strand is paired with a T from the converted strand (product 1) or G from the protected strand is paired with A from the converted strand (product 2). ii) The conversion-induced mismatches (C/T and G/A) are excluded from all filters. iii) Soft-clipping below a certain length is allowed at the 5’ end (default is 3bp, adjustable via command-line options). iv) Alignment score filter is turned off for the converted strands, though mapping quality filter is still applied. To calculate residual SNV frequencies by mutation contexts in NA12878, mutations detected in product 2 were reverse complemented (e.g., C>T became G>A) and combined with product 1 mutations. For the denominators (number of A,C,G,T bases) in the residual SNV frequencies, we complemented the bases detected in product 2 (e.g., C became G and G became C) and combined with the bases detected in product 1.

## Data availability

Methyl-CODEC sequencing data generated in this study will be available from dbGaP. EM-seq and WGBS NA12878 were downloaded from BioProject database (https://www.ncbi.nlm.nih.gov/bioproject/) under accession number PRJNA591788. Whole-genome sequencing (WGS) data for the HCT116 cell line was obtained from the BioProject database under accession number PRJNA875498. The HG38 CpG island coordinates are available at UCSC’s goldenpath service (http://hgdownload.cse.ucsc.edu/goldenpath/hg38/database/cpgIslandExt.txt.gz)

## Code availability

Methyl-CODEC analysis software and code to reproduce the results in the paper is available at https://github.com/broadinstitute/Methyl-CODEC, which is a fork from CODECsuite https://github.com/broadinstitute/CODECsuite.Bismark (v0.24.0) is available at https://github.com/FelixKrueger/Bismark. Picard is available at https://broadinstitute.github.io/picard/

## Acknowledgements

The authors acknowledge the Gerstner Family Foundation for its generous support. G.M.M. and V.A.A receive funding support from NIH National Cancer Institute (2R01CA221874-04).This study used the NA12787 cell line sample from the NINDS Repository.

## Author Contributions

R.Liu developed and performed the analyses. F.D. collected DNA samples and performed the experiments. V.A.A and G.M.M jointly designed and supervised the study. R.Liu, F.D., V.A.A and G.M.M developed the method and wrote the manuscript. L.W. contributed to the figures. R.Li contributed to the experiments, sequencing submissions and manuscript editing. J.H.B contributed to the adapter design and initial experiments. A.N. contributed to selecting dCTP variants. All authors interpreted the data, and all authors read, edited and approved the manuscript.

## Competing interests

A patent application has been filed on the methods disclosed in this manuscript. V.A.A., J.H.B., R.L. and G.M.M. are co-inventors on a patent application (PCT/US2021/062966) covering CODEC. V.A.A. and G.M.M. are co-inventors on a patent application covering the MAESTRO MRD test (US 2023/0203568) licensed to Exact Sciences which was not involved in this study. V.A.A. receives sponsored research funding from Exact Sciences and is a co-founder and advisor to Amplifyer Bio, which was not involved in this study.

## Notes

https://github.com/broadinstitute/Methyl-CODEC

### Competing Interest Statement

The authors have declared no competing interest.

