## Supplementary Fig. 1 for "Methyl-CODEC enables simultaneous methylation and duplex sequencing"

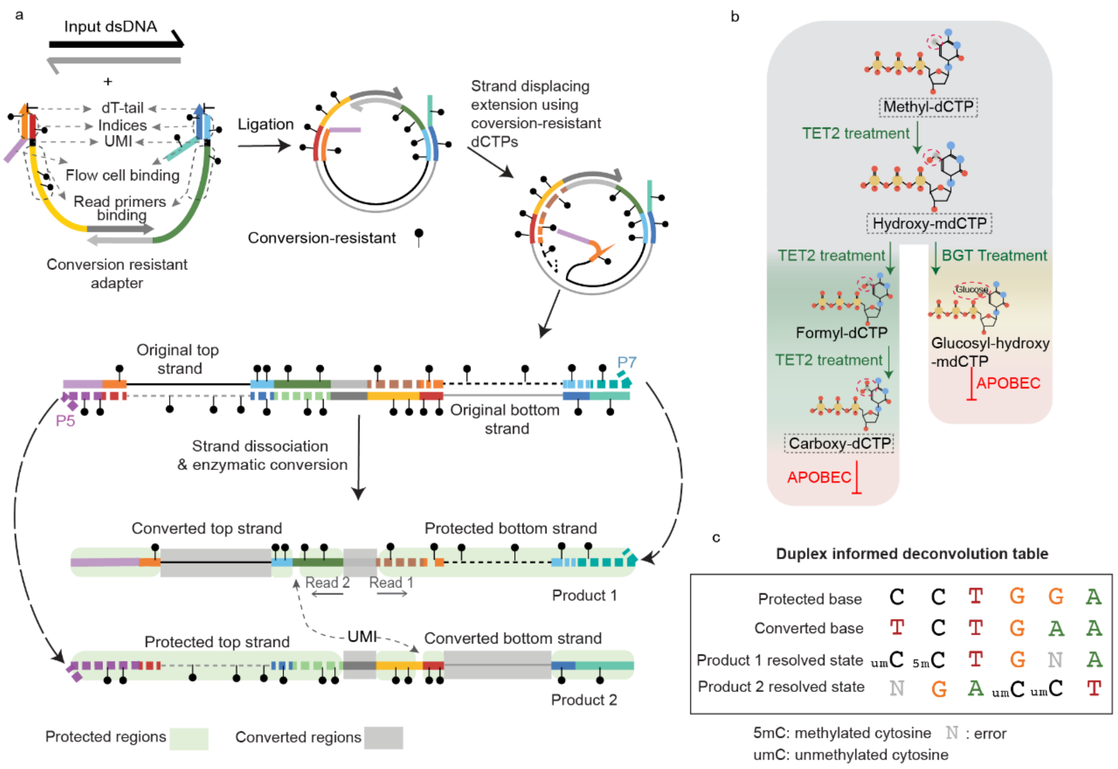


**Supplementary Fig. 1: Additional information for Methyl-CODEC workflow.**

**(a)** Detailed adapter and Methyl-CODEC product structures; The adapter quadruplex, which is resistant to conversion in its functional regions (circle with a dashed outline), is used for ligation, resulting in a loop-shaped ligated DNA. The strand-displacing extension is performed using Phi29 polymerase with conversion-resistant dCTP, instead of regular dCTP, producing an annealed DNA of two super strands. Each super strand consists of the original sequence (solid line) concatenated with the extended protected sequence (dashed line). The former retains the methylation pattern of the input DNA, while the latter remains intact during enzymatic methyl-conversion. Following strand dissociation and enzymatic methyl-conversion, each super strand on its own is a Methyl-CODEC product and allows for simultaneous methylation and duplex sequencing.

**(b)** Conversion-resistant dCTP variants used in Methyl-CODEC; TET2 and BGT enzymes convert methyl-dCTP and hydroxy-methyl-dCTP into carboxy-dCTP and glucosyl-hydroxy-methyl-dCTP which are resistant to conversion due to structural modifications that prevent recognition by the APOBEC enzyme. In addition to methyl-dCTP, hydroxy-methyl-dCTP, and carboxy-dCTP, we also tested propargylamino-dCTP, a synthetic variant not naturally occurring in the body. Methyl-CODEC results indicates that propargylamino-dCTP is also a conversion-resistant dCTP variant.

c) Duplex deconvolution table: the bases A, G, T, unmethylated C and methylated C are determined by comparing the complementary bases from the two strands, according to whether the read pair corresponds to product 1 vs product 2. Cases not addressed by the table are considered as errors and hence suppressed.
