## Supplementary Fig. 2 for "Methyl-CODEC enables simultaneous methylation and duplex sequencing"

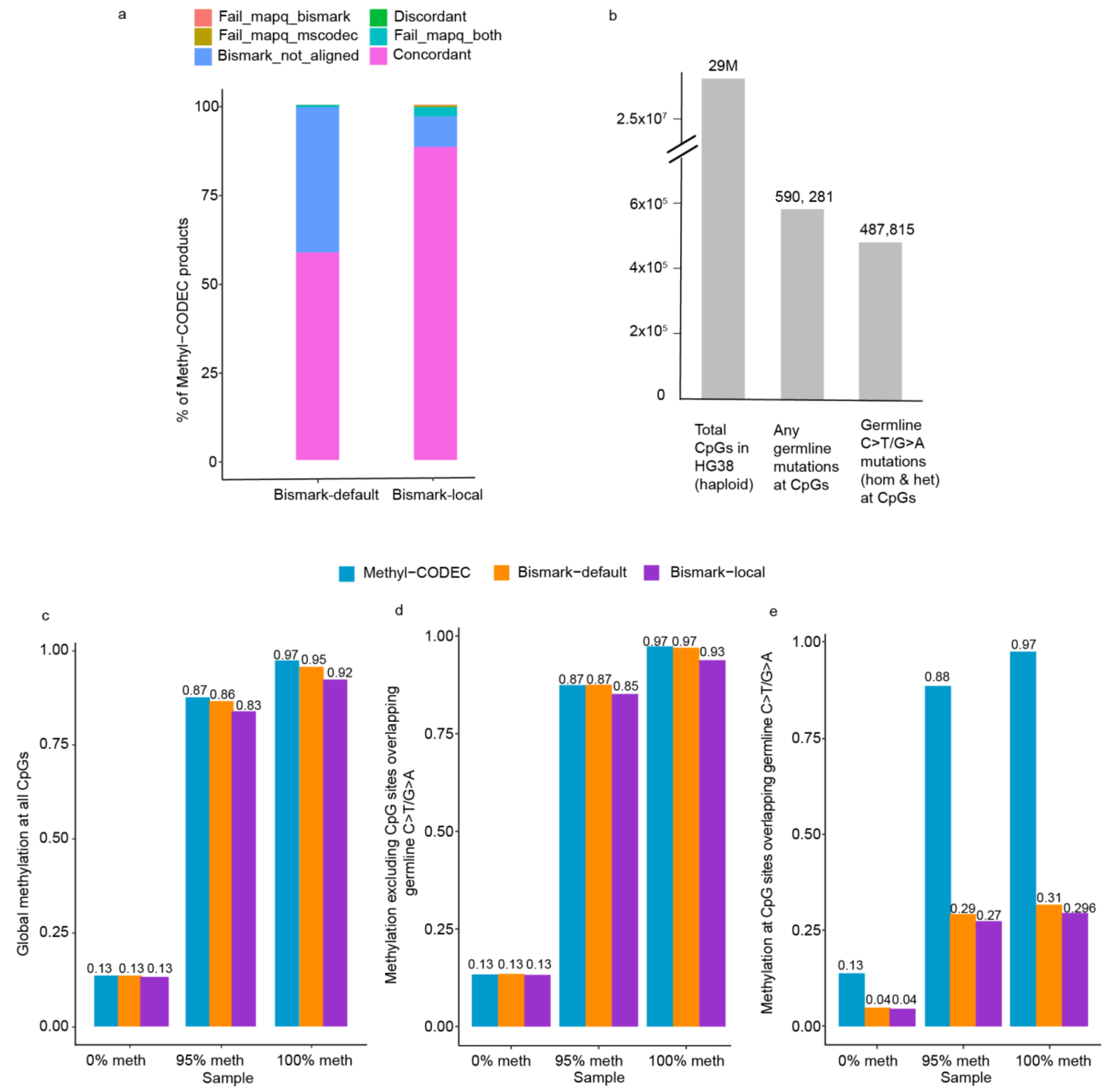


**Supplementary Fig. 2: Additional advantages of preserving original DNA for methylation sequencing.** **(a)** Comparing NA12878 Methyl-CODEC alignments vs Bismark alignments with two different parameter modes (default vs local). For Bismark alignment, we simulated paired-end methyl-converted reads by in-silico converting the protected strand from Methyl-CODEC products. concordant: concordant alignments between Methyl-CODEC and Bismark, inferred by distances of the two alignments less than 500 bp; bismark_not_align: reads were aligned by Methyl-CODEC but not by Bismark; discordant: reads aligned to different regions in the genome by Methyl-CODEC and Bismark (at least 500 bp apart from each other); fail_mapq_bismark: reads below mapping quality cutoff for Bismark alignment but satisfying mapping quality cutoff for Methyl-CODEC alignment; fail_mapq_mscodec: reads below mapping quality cutoff for Methyl-CODEC alignment but satisfying mapping quality cutoff for Bismark alignment; fail_mapq_both:

reads below mapping quality cutoff for both Methyl-CODEC and Bismark alignments. Mapping quality cutoff was determined by the highest mapping quality of the respective aligner (Bowtie2 for Bismark and BWA for Methyl-CODEC) and was required to be a multiple of 10 (Bismark: 40, Methyl-CODEC: 60).

**(b)** Bar plot of numbers of total CpGs in HG38, CpGs overlapping any HCT116 germline mutations (including Indels and SNVs) and CpGs overlapping only germline C>T and G>A mutations.

**(c-e)** Comparison of methylation calling results from HCT116 Methyl-CODEC vs simulated EM-seq WGMS. Paired-end EM-seq WGMS was simulated by in-silico methyl-converting the protected strands of the Methyl-CODEC samples and then aligned by Bismark (default and local modes) (c) Average sample level CpG methylation for unmethylated control, methylated control and 1 to 19 ratio mixing of the two. (d) Average CpG methylation per sample when excluding CpGs overlapping germline C>T and G>A mutations for the unmethylated control, methylated control and 1 to 19 ratio mixing of the two. (e) Average methylation at CpGs overlapping germline C>T and G>A mutations for the unmethylated control, methylated control and 1 to 19 ratio mixing of the two.
