## Supplementary Fig. 3 for "Methyl-CODEC enables simultaneous methylation and duplex sequencing"

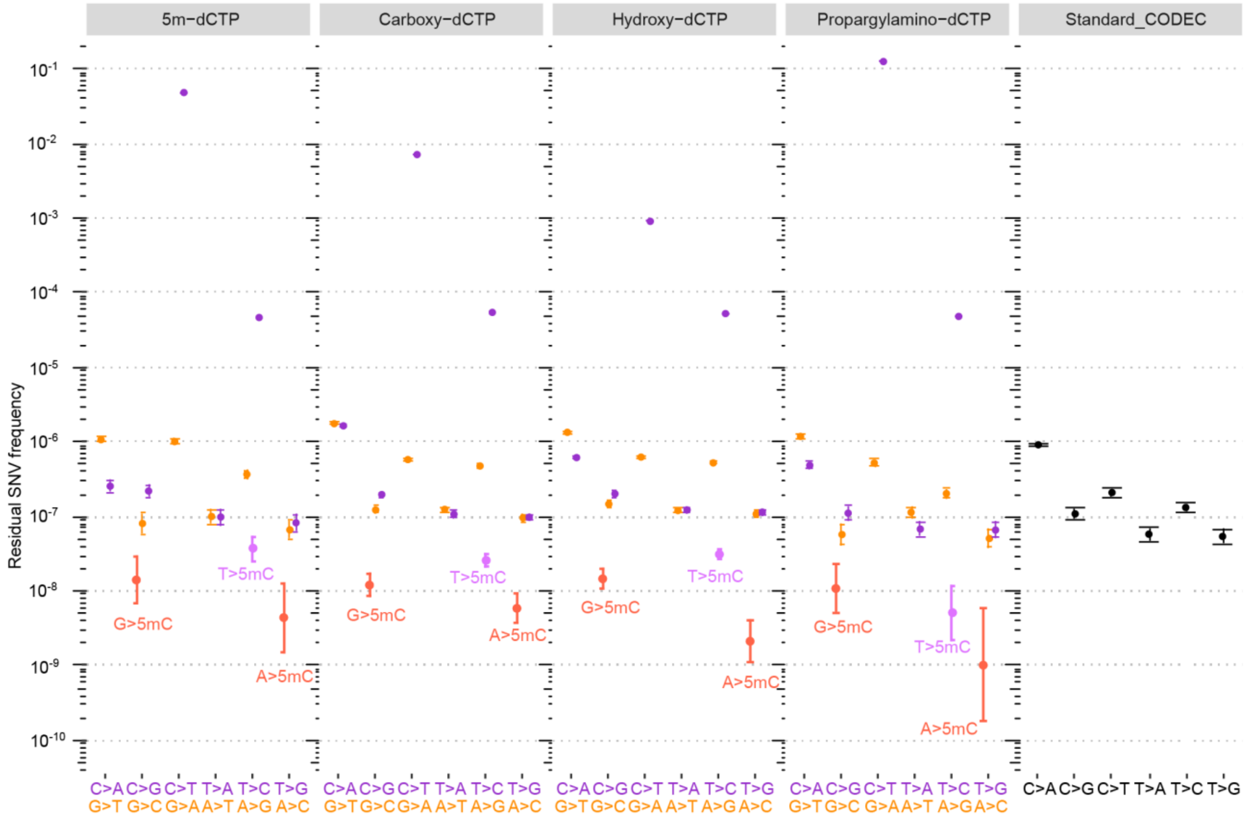


**Supplementary Fig. 3:** Scatter plot comparing residual SNV frequency of standard CODEC and 4 different versions of Methyl-CODEC (using 4 different conversion-resistant dCTP variants) in NA12878, all using a commercial end-repair protocol. Orange and purple colors are used to indicate mutations with C&T and G&A as the reference bases, respectively. Mutations that produce unmethylated cytosines were colored by orange and purple as well. Whereas mutations that produce methylated cytosines (5mC) are distinguished by slightly different colors and are labeled next to the corresponding dots.
